# A pleiotropic chemoreceptor facilitates the functional coupling of pheromone production and perception

**DOI:** 10.1101/2022.01.10.475668

**Authors:** Cassondra Vernier, Kathleen M. Zelle, Nicole Leitner, Xitong Liang, Sean Halloran, Jocelyn G. Millar, Yehuda Ben-Shahar

**Affiliations:** Department of Biology, Washington University in Saint Louis, 1 Brookings Drive, Saint Louis, MO 63130, USA; Carl R. Woese Institute for Genomic Biology, University of Illinois, 1206 W. Gregory Dr., Urbana, IL 61801, USA; Department of Neuroscience, Washington University School of Medicine, 660 South Euclid Avenue, St. Louis, MO 63110, USA; Department of Entomology, University of California, Riverside, 900 University Avenue, Riverside, CA 92521, USA

**Keywords:** *Drosophila melanogaster*, Vinegar fly, Fruit fly, Cuticular hydrocarbons, *Gr8a*, Oenocyte

## Abstract

Optimal mating decisions depend on the robust coupling of signal production and perception because independent changes in either could carry a fitness cost. However, since the perception and production of mating signals are often mediated by different tissues and cell types, the mechanisms that drive and maintain their coupling remain unknown for most animal species. Here, we show that in *Drosophila,* sensory perception and production of an inhibitory mating pheromone are co-regulated by *Gr8a,* a member of the *Gustatory receptor* gene family. Specifically, we found that the pleiotropic action of *Gr8a* independently regulates the perception of pheromones by the chemosensory systems of males and females, as well as their production in the fat body and oenocytes of males. These findings provide a relatively simple molecular explanation for how pleiotropic receptors maintain robust mating signaling systems at the population and species levels.

## INTRODUCTION

The majority of sexually-reproducing animals use intricate mating signaling systems, which rely on a robust physiological coupling between the production and perception of species-specific signals since any independent changes in either the signal or the capacity to sense it would carry a fitness cost (Boake, 1991; Brooks et al., 2005; Hoy et al., 1977; Shaw et al., 2011; Shaw and Lesnick, 2009; Steiger et al., 2011; Sweigart, 2010; Symonds and Elgar, 2008; Wyatt, 2014). Previously published theoretical models have postulated that the maintenance of robust coupling between the production and perception of mating signals is driven by strong genetic linkage between the cellular and physiological processes that regulate mating-signal production and its perception, or alternatively, via the action of pleiotropic genes that control both processes (Boake, 1991; Butlin and Ritchie, 1989; Butlin and Trickett, 1997; Shaw *et al*., 2011; Shaw and Lesnick, 2009). Consequently, both mechanisms provide plausible explanations for how mating-signaling systems could remain stable and reliable at the population level while still retaining their capacity for future diversification, as necessitated for speciation (Chebib and Guillaume, 2021; Hoy *et al*., 1977; Kirkpatrick and Hall, 2004; Lande, 1980; Shaw *et al*., 2011; Shaw and Lesnick, 2009; Wiley et al., 2012).

Empirical data in support of the contribution of gene-linkage or pleiotropy to the maintenance of coupling between mating signal production and perception at the population level are rare (Chebib and Guillaume, 2021; Hoy *et al*., 1977; Shaw *et al*., 2011; Shaw and Lesnick, 2009; Wiley *et al*., 2012). Additionally, the complex characteristics of mating behaviors, and the species-specific signals that drive them, present a major barrier for identifying the actual molecular mechanisms and candidate pleiotropic genes that support the coupling between the production and perception of specific mating signals (Chenoweth and Blows, 2006; Singh and Shaw, 2012). How the functional coupling of the physiological processes responsible for the production and perception of mating signals remains robust is particularly puzzling since their perception is mediated by the peripheral sensory nervous system, while their production is restricted to specialized, non-neuronal pheromone producing cells (Chung and Carroll, 2015; Chung et al., 2014; McKinney et al., 2015). Notwithstanding, a previous *Drosophila* study has implied that the gene *desat1,* which encodes a fatty acid desaturase, directly contributes to both the perception and production of pheromones (Bousquet et al., 2012). However, subsequent studies have shown that *desat1* expression is enriched in central neurons, and that the effect of *desat1* mutations on the behavioral response to pheromones is not likely to be directly mediated via the modulation of pheromone perception by sensory neurons (Billeter et al., 2009). Furthermore, the effects of *desat1* mutations on the overall CHC profiles of both males and females are broad and lack specificity (Labeur et al., 2002). Together, these data suggest that *desat1* is not likely to act as a pleiotropic factor that directly couples the production and perception of mating pheromones in *Drosophila.* Consequently, the molecular identities of genes that may mediate the genetic and functional linkage between the production of insect mating pheromones by the coenocytes, and their perception by the chemosensory system, remained unknown.

Here we show that some pheromone-driven mating behaviors in *Drosophila* depend on the pleiotropic action of *Gr8a,* a member of the *Gustatory receptor* gene family (Lee et al., 2012; Shim et al., 2015), which contributes to both the perception of inhibitory mating signals in pheromone-sensing neurons, and independently, to the production of inhibitory mating pheromones in non-neuronal abdominal pheromone-producing oenocytes. Together, these data provide a relatively simple molecular explanation for how genetic linkage could maintain functional coupling between the independent cellular and physiological processes that drive pheromone perception and production.

## RESULTS

### Some gustatory-like receptors exhibit enriched expression in abdominal tissues

Similar to other insect species, *Drosophila* cuticular hydrocarbons (CHCs), or long-chain fatty acids synthesized by the fat body and oenocytes (Billeter *et al*., 2009; Gutierrez et al., 2007), provide a hydrophobic desiccation barrier, as well as play an important role as pheromones in regulating diverse behaviors, including mating (Blomquist and Bagnères, 2010; Chung and Carroll, 2015; Ferveur, 2005; McKinney *et al*., 2015). Specifically, complex blends of CHCs are often utilized by insects to communicate sex identity and female mating status, as well as to define the behavioral reproductive boundaries between closely related species (Ben-Shahar, 2015; Billeter *et al*., 2009; Chung and Carroll, 2015; Chung *et al*., 2014; Dweck et al., 2015; Lu et al., 2012; Lu et al., 2014; Yew and Chung, 2015).

While some of the genes and pathways that contribute to CHC synthesis in *Drosophila* are known (Blomquist and Bagnères, 2010; Chung *et al.*, 2014; Ferveur, 2005; Howard and Blomquist, 2005; McKinney *et al*., 2015), the molecular identities of most CHC receptors remain unknown. Current models stipulate that the perception of volatile CHCs is mediated by olfactory sensory neurons (ORNs) located in the antennae and maxillary palps, while less volatile CHCs are sensed by specialized gustatory-like receptor neurons (GRNs) in the appendages (legs and wings), female genitalia, and the proboscis (Benton et al., 2007; Clowney et al., 2015; Datta et al., 2008; Koh et al., 2014; Kurtovic et al., 2007; Lebreton et al., 2014; Lu *et al*., 2012; Lu *et al*., 2014; Pikielny, 2012; Thistle et al., 2012; Toda et al., 2012; van der Goes van Naters and Carlson, 2007; Vijayan et al., 2014).

Consequently, we chose to examine members of the *Gustatory receptor (Gr)* gene family as candidate pleiotropic genes that might contribute to both the perception and production of pheromonal mating signals in *Drosophila.* Because several family members have already been implicated in the detection of specific excitatory and inhibitory pheromones (Bray and Amrein, 2003; Miyamoto and Amrein, 2008; Moon et al., 2009; Watanabe et al., 2011), and the majority of genes that encode family members are already known to be enriched in GRNs (Clyne et al., 2000; Dunipace et al., 2001; Scott et al., 2001; Wang et al., 2004), we reasoned that any pleiotropic *Gr* genes should be also expressed in the abdominal oenocytes (Billeter *et al*., 2009). We tested this by using an RT-PCR screen, which revealed that 24 out of the 60 members of the *Gr* family are expressed in abdominal tissues of adult *Drosophila* (Table 1). This suggests that at least some *Gr* genes may contribute to both the perception and production of mating signals in *Drosophila.*

**Table 1.** Candidate *Gr* genes expressed in male and/or female abdomens. Plus and minus signs indicate whether RT-PCR products were detected. Only genes with positive PCR products in at least one sex are shown.

| **Gene** | **Male** | **Female** |
| --- | --- | --- |
| *Gr2a* | - | + |
| *Gr8a* | + | - |
| *Gr10a* | + | + |
| *Gr21a* | - | + |
| *Gr22a* | + | - |
| *Gr22e* | + | + |
| *Gr36c* | + | - |
| *Gr58c* | + | + |
| *Gr59a* | + | + |
| *Gr59b* | + | + |
| *Gr63a* | + | - |
| *Gr64a* | + | - |
| *Gr64b* | + | + |
| *Gr64c* | + | + |
| *Gr64d* | + | - |
| *Gr66a* | + | + |
| *Gr89a* | + | + |
| *Gr93a* | - | + |
| *Gr93d* | + | + |
| *Gr97a* | + | + |
| *Gr98a* | + | + |
| *Gr98b* | - | + |
| *Gr98c* | + | + |
| *Gr98d* | + | + |

### Gr8a is a chemosensory receptor with sexually dimorphic expression in abdominal cells

Although several members of the *Gr* gene family, including *Gr68a, Gr32a, Gr66a, Gr39a,* and *Gr33a,* were previously linked to the sensory perception of mating pheromones (Bray and Amrein, 2003; Lacaille et al., 2007; Miyamoto and Amrein, 2008; Moon *et al*., 2009; Watanabe *et al*., 2011), none of these candidate genes were identified in our initial RT-PCR screen for *Gr* genes expressed in abdominal tissues of either males or females (Table 1). However, *Gr8a,* which was indicated by our screen as being a male-specific abdomen-enriched receptor (Table 1) (Park and Kwon, 2011), was previously shown to play a role in the chemosensation of the non-proteinogenic amino acid L-Canavanine (Lee et al., 2012; Shim et al., 2015). Because our initial expression screen was based on whole-abdomen RNAs, we next used a GAL4 transgenic driver to determine which abdominal cells express *Gr8a.* We found that, as was previously reported (Lee *et al*., 2012), *Gr8a* is expressed in 14-16 GRNs in the proboscis (Figure 1A-B), as well as in two paired GRNs in the pretarsus of the prothoracic legs in males (Figure 1C) and females (Figure 1D). We also observed *Gr8a* expression in abdominal oenocyte-like cells in males (Figure 1E) but not females (Figure 1F). The male-biased expression in the abdomen was further supported by qRT-PCR analysis (Figure 1G). These data further indicate that in addition to its chemosensory functions, *Gr8a* may also contribute to oenocyte physiology.

**Figure 1.**
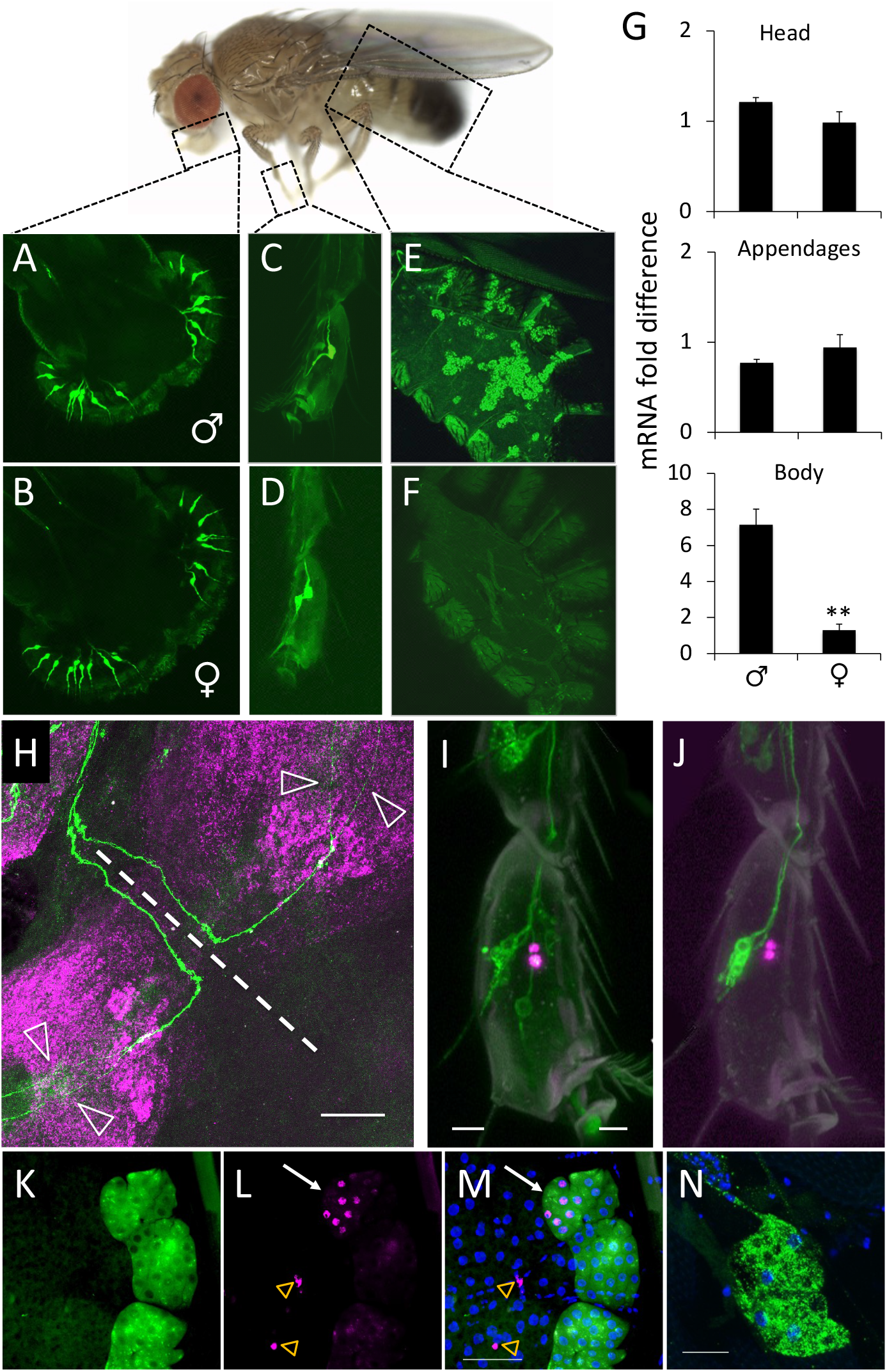
*Gr8a* is a sexually dimorphic chemosensory receptor. **(A-F)** *Gr8a* is expressed in the proboscis (A-B) and prothoracic legs (C-D) of both males (top) and females (bottom), but is only expressed in the abdomen of males (E-F). Cells labeled by *Gr8a-*GAL4. **(G)** *Gr8a* has sexually dimorphic mRNA expression in the bodies of flies. Relative mRNA levels were measured by real-time quantitative RT-PCR. **, p<0.01 Mann Whitney Rank Sum Test, n=3/group. **(H-J)** *Gr8a*-expressing GRNs represent a distinct subclass of pheromone sensing neurons. (H) Axonal projection patterns in the T1 VNC neuromere in a *Gr8a*-GAL4>UAS-CD8::GFP male (green). Arrowheads, individual axons; dashed line, midline of the VNC. Magenta, neuropil marker (nc82). (I) Confocal z-stack of a male *fruP1*-LexA>LexAop-myrGFP (green); *Gr8a*-GAL4>UAS-Red-Stinger (magenta) prothoracic leg. (J) Confocal z-stack of a male *ppk23-* LexA>LexAop-CD8::GFP (green); *Gr8a*-GAL4>UAS-Red-Stinger (magenta) prothoracic leg. **(K-M)** *Gr8a* is expressed in oenocytes and other abdominal cells. Confocal z-stack images of oenocytes in a *Gr8a*-GAL4>UAS-CD8::GFP; *desat1>luciferase* male: (K) *desat1* (green); (L) *Gr8a* (magenta); (M) Merge. Blue, DAPI. White arrow, expression of *Gr8a* in oenocytes; yellow arrows, expression of *Gr8a* in other abdominal cells. **(N)** GR8A protein is enriched in abdominal cells. Confocal z-stack of a GFP-tagged *Gr8a* allele in male abdominal cells; green, anti-GFP; blue, DAPI. Scale bars = 50μm.

To further examine the spatial expression of *Gr8a* in males, we used a membrane bound GFP reporter to trace the axonal projection patterns of *Gr8a*-expressing GRNs in the prothoracic legs. We found that in contrast to the primary, sexually dimorphic *ppk23-* expressing pheromone-sensing GRNs (Lu *et al*., 2012; Lu *et al*., 2014; Thistle *et al*., 2012; Toda *et al*., 2012), the axons of tarsal *Gr8a-*expressing neurons ascend to the brain and do not cross the midline of the ventral nerve cord (VNC) in males (Figure 1H). Likewise, we found that *Gr8a*-expressing GRNs do not co-express the sex determination factor *fru* (Figure 1I) or the ion channel *ppk23* (Figure 1J), which are were previously assumed to be expressed in all pheromone-sensing GRNs in the fly appendages. These data indicate that *Gr8a*-expressing GRNs in the prothoracic tarsal segments possibly represent a distinct subclass of pheromone-sensing GRNs.

In the male abdomen, we found that *Gr8a* is co-expressed with the oenocyte specific *desat1* driver (Billeter et al., 2009), as well as possibly in *desat1*-negative fat-body-like cells (Figure 1K-M). To better understand how Gr8a might function in non-neuronal oenocytes, we next characterized the subcellular localization of the native *Gr8a* protein in abdominal tissues, by using CRISPR/Cas9 genome editing to generate an endogenous GFP-tagged allele of *Gr8a.* Subsequently, immunohistochemical staining of abdominal tissues from *Gr8a*-GFP males with an anti-GFP antibody revealed that the receptor protein is enriched in vacuolar membranes in some oenocyte clusters (Figure 1N). Together, these data indicate that in addition to its possible role in the perception of L-Canavanine, *Gr8a* also contributes to the perception, and possibly production, of mating pheromones in the male.

### Gr8a activity contributes to mating decisions in females

We next hypothesized that if *Gr8a* is a pleiotropic gene that independently contributes to the production of a mating pheromone in males, and its chemosensory perception in females, then the knockdown of *Gr8a* in either males or females should have similar effects on female mating behavior. Therefore, we first investigated whether *Gr8a,* and the GRNs that express it, are required for sensory functions associated with female mate choice by using single-pair courtship assays (Lu *et al*., 2012; Lu *et al*., 2014). We found that blocking neuronal transmission in female *Gr8a*-expressing GRNs by the transgenic expression of tetanus toxin (TNT) shortens copulation latency relative to wild-type females, when courted by wild-type males (Figure 2A). Similarly, homozygous (Figure 2B) and hemizygous (Figure 2C) *Gr8a*-null females exhibited shorter copulation latencies when courted by wild type males, which can be rescued by driving the expression of the *Gr8a* cDNA by *Gr8a*-GAL4 (Figure 2D). In contrast, genetic manipulations of *Gr8a* did not affect male courtship behavior as measured by courtship latency and index towards wild-type females (Supplemental Figure 1). These data suggest that *Gr8a* is required for regulating female mating receptivity via the chemosensory detection of male-borne inhibitory mating pheromones.

**Figure 2.**
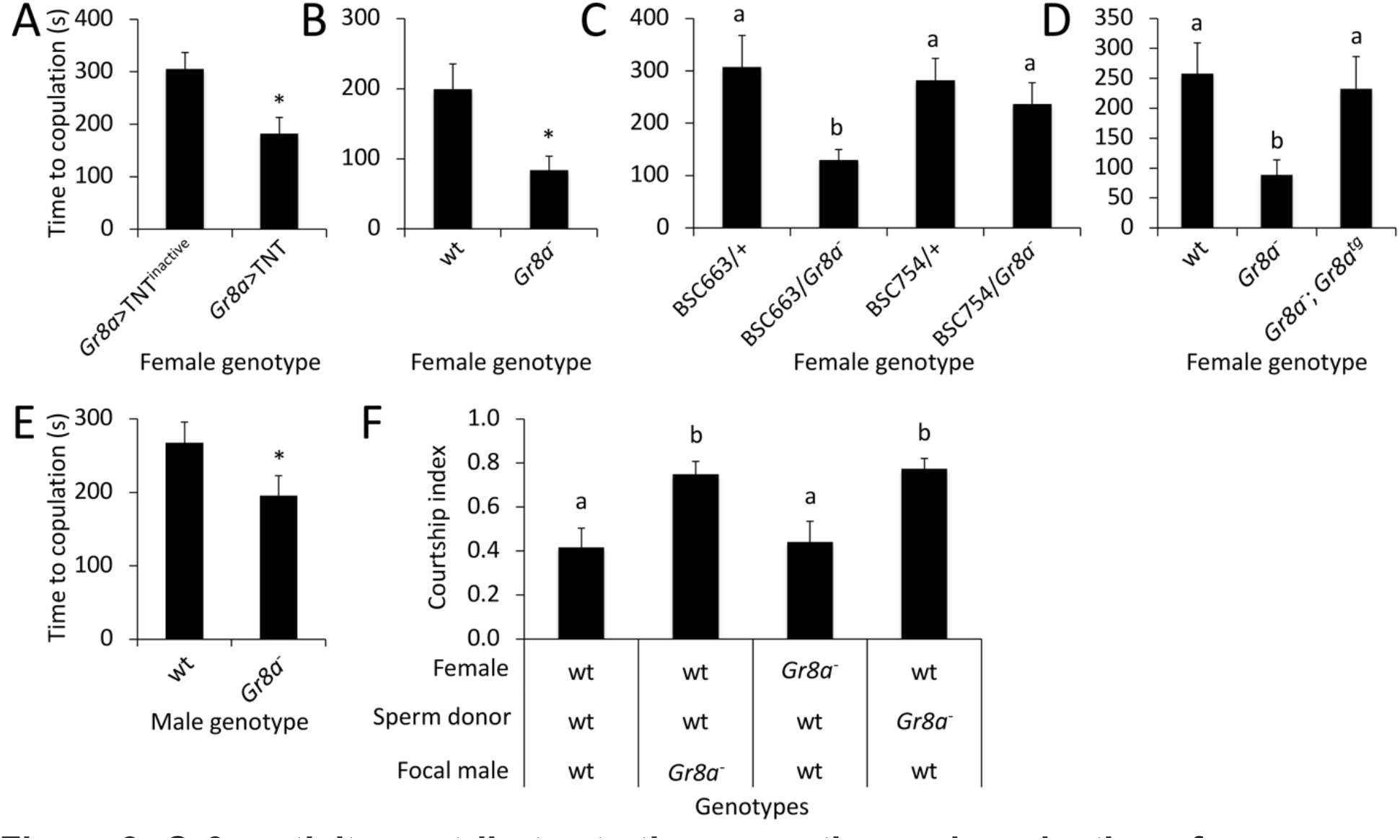
*Gr8a* activity contributes to the perception and production of an inhibitory signal associated with mating decisions in males and females. **(A)** Blocking neural activity in female *Gr8a*-expressing sensory neurons (*Gr8a*>TNT) shortens copulation latency relative to wild-type controls (*Gr8a*>TNT^inactive^). **(B-C)** Homozygous (B) or hemizygous (C) *Gr8a* null females show shortened copulation latency relative to wild-type controls. Df(1)BSC663 is a deficiency that covers the *Gr8a* locus. Df(1)BSC754 was used as a control. **(D)** Expression of *Gr8a* cDNA with the *Gr8a* promoter (Gr8a-;Gr8a^tg^) rescues the copulation latency phenotype in *Gr8a* mutant females. **(E)** Wild-type females exhibit shortened copulation latency when courted by *Gr8a* mutant males relative to wild-type males. **(F)** *Gr8a* mutant males do not recognize the mating status of females, and have a reduced transfer of inhibitory mating pheromones during copulations. Female, female genotype; Sperm donor, genotype of males mated first with focal females; Focal male, genotypes of experimental males presented with mated females. Different letters above bars indicate statistically significant Tukey’s HSD *post hoc* contrasts between groups. Panels C, D, and F: p<0.05 ANOVA, n>15/group. Panels A, B, E: *, p<0.05, Mann Whitney Rank Sum Test, n>15/group. All assays performed under red light conditions.

Because *Gr8a* expression is specifically enriched in male oenocytes (Figure 1K-M), we next tested the hypothesis that *Gr8a* also plays a role in the production and/or release of inhibitory mating signals by males. We found that wild-type virgin females exhibited shorter copulation latencies towards *Gr8a* mutant males relative to wild-type controls (Figure 2E). These data indicate that the *Gr8a* mutant males produce and/or release lower levels of inhibitory mating pheromones relative to wild type controls. Together, these behavioral studies suggest that *Gr8a* is a pleiotropic gene that regulates both the production of an inhibitory mating signal in the male oenocytes, and its perception by the chemosensory system in females.

### Gr8a regulates the copulatory transfer, and the post-mating perception, of inhibitory pheromones by males

Mating decisions in *D. melanogaster* rely on a balance between excitatory and inhibitory drives (Billeter *et al*., 2009; Clowney *et al*., 2015; Kallman et al., 2015; Krupp et al., 2008; Laturney and Billeter, 2016). Therefore, male-borne inhibitory signals may help females optimize mate choices by delaying their decision to copulate with specific males. Additionally, previous studies showed that, in order to increase their fitness, *Drosophila* males transfer inhibitory mating pheromones to females during copulation, which subsequently lowers the overall attractiveness of mated females to other males (Averhoff and Richardson, 1974; Datta *et al*., 2008; Jin et al., 2008; Kurtovic *et al*., 2007; Miyamoto and Amrein, 2008; Yang et al., 2009). We found that *Gr8a* mutant males were more likely to court mated females than wild-type controls (Figure 2F), suggesting that *Gr8a* is also required in males for the sensory recognition of the inhibitory signals that label the post-mating status of females. We also found that wild-type males failed to recognize the mating status of wild-type females that were previously mated with *Gr8a* mutant males (Figure 2F). These data indicate that *Gr8a* is also important for the production of inhibitory pheromones that are transferred from males to females during copulation. Together, these findings suggest that *Gr8a* is responsible for the production and perception of transferrable inhibitory mating signals that advertise post-mating status in females. The simplest overall interpretation of these data is that *Gr8a* is a pleiotropic factor, which independently contributes to the production/ transfer of male inhibitory mating pheromones, as well as their sensory perception in both males and females.

### Gr8a contributes to quantitative and qualitative attributes of the pheromone profiles of males and mated females

Because our data indicate that the *Gr8a* mutation has a dramatic effect on the copulation latency of mated females and the ability of males to detect the mating status of females, we hypothesized that *Gr8a* is contributing to the production and/or transfer of an inhibitory pheromone in males. Therefore, we next examined whether the *Gr8a* mutation has a direct effect on qualitative and quantitative aspects of male and mated-female CHC profiles. We found that the overall CHC profile of *Gr8a* mutant males is both qualitatively (Figure 3A) and quantitatively different from that of wild-type males (Figure 3B-C and Table 2). In particular, the *Gr8a* mutation affects the levels of several alkenes and methyl-branched alkanes, which have been implicated in mate choice behaviors in diverse *Drosophila* species (Billeter *et al.*, 2009; Billeter and Levine, 2013; Chung and Carroll, 2015; Chung *et al*., 2014; Dyer et al., 2014; Shirangi et al., 2009).

**Figure 3.**
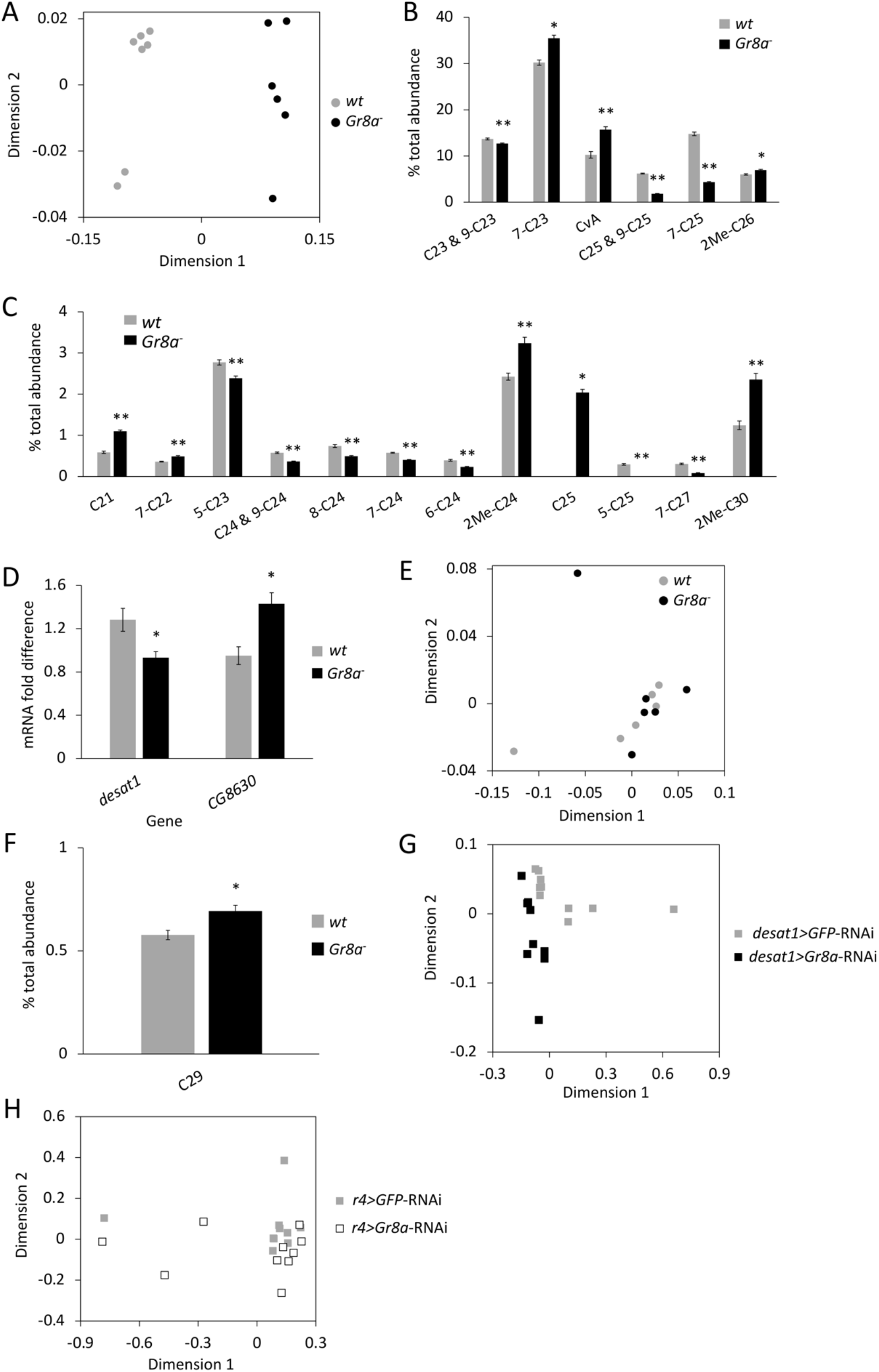
The *Gr8a* mutation affects the pheromone profiles of males and mated females. **(A)** Wild-type (wt) and *Gr8a* mutant (Gr8a^-^) males differ in CHC profile. p<0.001, Permutation MANOVA. **(B-C)** The *Gr8a* mutation affects the levels of individual CHCs in males. (B) CHCs found at high proportions in males. (C) CHCs found at low proportions in males. Only affected CHCs are shown. See Table 2 for the complete list. *, p<0.05, **, p<0.001, Student’s t-test or Mann Whitney Rank Sum Test, n=6 (Gr8a-) or 7 (wt). **(D)** The *Gr8a* mutation affects the expression level of several desaturase genes. Only affected genes are shown. See Table 3 for the complete list. *, p<0.05, Student’s t-test, n=4/group. **(E)** Females mated with wild-type or *Gr8a* mutant males do not differ in CHC profile. p=0.570, Permutation MANOVA. **(F)** Nonacosane (C29) differs between females mated with wild-type and *Gr8a* mutant males. See Table 4 for complete list of mated-female CHCs. *, p<0.05, Student’s t-test, n=6/group. **(G)** Control *(desat1 > GFP-RNAi)* and oenocyte-specific *Gr8a* knockdown *(desat1 > Gr8a-* RNAi) males differ in CHC profile. p<0.001, Permutation MANOVA. **(H)** Control *(r4 > GFP*-RNAi) and fat body-specific *Gr8a* knockdown (*r4 > Gr8a*-RNAi) males do not differ in CHC profile. p = 0.298, Permutation MANOVA. Panels A, E, G, and H depicted as Nonmetric Multidimensional Scaling (NMDS) plots with Bray-Curtis dissimilarity.

**Table 2.** Male CHCs. Retention time (R.T.), compound, percent total (% total), and p-value (Student’s t-test or Mann Whitney Rank Sum Test) of each compound as part of the total pheromonal bouquet for wild-type (wt) and *Gr8a* mutant (*Gr8a^-^*) males.

| R.T. | Compound | wt % total | *Gr8a*<sup>-</sup> % total | p value |
| --- | --- | --- | --- | --- |
| 12.31 | C<sub>21</sub> | 0.589 | 1.102 | <0.001 |
| 13.24 | Unknown | 0.071 | 0.191 | <0.001 |
| 14.2 | C<sub>22</sub> | 0.893 | 0.949 | 0.339 |
| 14.34 | 7-C<sub>22</sub> | 0.362 | 0.490 | <0.001 |
| 15.25 | Unknown | 0.128 | 0.226 | <0.001 |
| 16.22 | C<sub>23</sub> & 9-C<sub>23</sub> | 13.682 | 12.671 | <0.001 |
| 16.4 | 7-C<sub>23</sub> | 30.201 | 35.478 | 0.002 |
| 16.53 | 5-C<sub>23</sub> | 2.772 | 2.389 | <0.001 |
| 16.71 | CvA | 10.240 | 15.700 | <0.001 |
| 18.03 | C<sub>24</sub> & 9-C<sub>24</sub> | 0.578 | 0.367 | <0.001 |
| 18.19 | 8-C<sub>24</sub> | 0.742 | 0.493 | <0.001 |
| 18.27 | 7-C<sub>24</sub> | 0.579 | 0.402 | <0.001 |
| 18.37 | 6-C<sub>24</sub> | 0.395 | 0.233 | <0.001 |
| 18.46 | 5-C<sub>24</sub> | 0.040 | 0.047 | 0.943 |
| 19.09 | 2Me-C<sub>24</sub> | 2.426 | 3.240 | <0.001 |
| 19.95 | C<sub>25</sub> | 0.000 | 2.038 | 0.001 |
| 20.02 | C<sub>25</sub> & 9-C<sub>25</sub> | 6.195 | 1.793 | <0.001 |
| 20.18 | 7-C<sub>25</sub> | 14.781 | 4.344 | <0.001 |
| 20.42 | 5-C<sub>25</sub> | 0.296 | 0.000 | <0.001 |
| 22.89 | 2Me-C<sub>26</sub> | 6.007 | 6.933 | 0.002 |
| 23.7 | C<sub>27</sub> | 1.127 | 0.719 | 0.052 |
| 23.94 | 7-C<sub>27</sub> | 0.308 | 0.083 | <0.001 |
| 26.46 | 2Me-C<sub>28</sub> | 4.992 | 5.775 | 0.078 |
| 27.25 | C<sub>29</sub> | 0.351 | 0.317 | 0.574 |
| 29.89 | 2Me-C<sub>30</sub> | 1.245 | 2.353 | <0.001 |

**Table 3.**
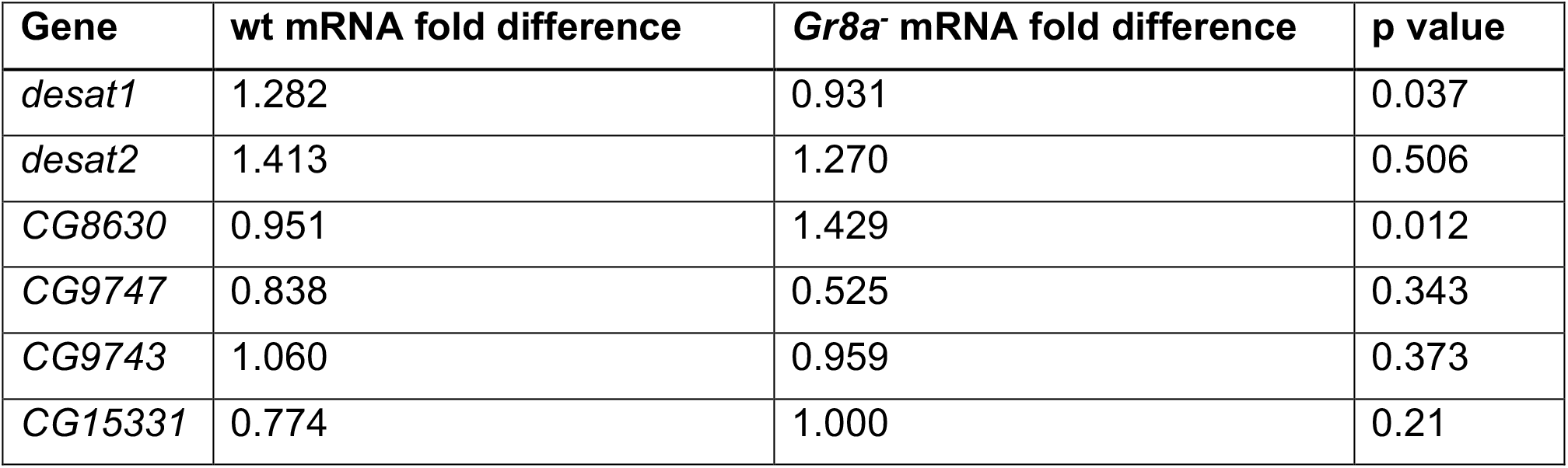
Desaturase gene expression. Relative mRNA expression of each desaturase gene for wild-type (wt) and *Gr8a* mutant *(Gr8a^-^)* males. Statistics via Student’s t-test.

| Gene | wt mRNA fold difference | *Gr8a*<sup>-</sup> mRNA fold difference | p value |
| --- | --- | --- | --- |
| *desat1* | 1.282 | 0.931 | 0.037 |
| *desat2* | 1.413 | 1.270 | 0.506 |
| *CG8630* | 0.951 | 1.429 | 0.012 |
| *CG9747* | 0.838 | 0.525 | 0.343 |
| *CG9743* | 1.060 | 0.959 | 0.373 |
| *CG15331* | 0.774 | 1.000 | 0.21 |

**Table 4.**
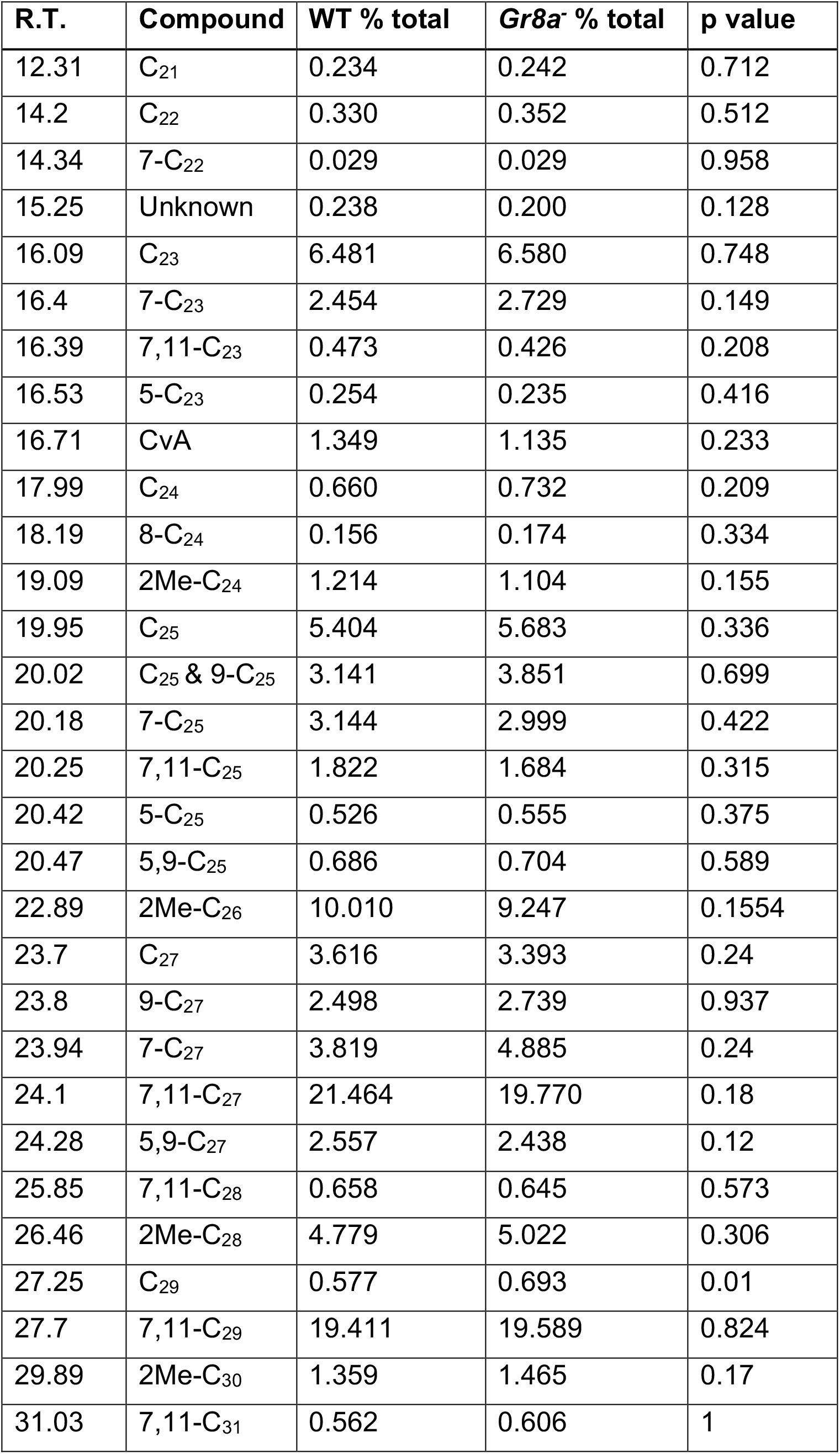
Mated-female CHCs. Retention time (R.T.), compound, percent total (% total), and p-value (Student’s t-test or Mann Whitney Rank Sum Test) of each compound as part of the total pheromonal bouquet for females mated with wild-type (wt) or *Gr8a* mutant *(Gr8a^-^)* males.

Although the exact mechanism by which *Gr8a* might be regulating the levels of specific CHCs remains unknown, we found that the expression levels of the desaturases *desat1* and *CG8630,* which play a role in the biosynthesis of alkenes (Chung and Carroll, 2015), are affected by the *Gr8a* mutation in the male abdomen (Figure 3D). We also found that the overall qualitative aspects of the CHC profiles of wild-type females were not affected by mating with either *Gr8a* mutant or wild-type males (Figure 3E). However, quantitative analyses of individual CHCs revealed that nonacosane (C29) is higher in females that mated with *Gr8a* mutant males relative to those that mated with wild-type males (Figure 3F). Together, these data suggest that *Gr8a* action in oenocytes contributes to the production of some cuticular alkenes and methyl-branched alkanes in males, which possibly function as inhibitory mating pheromones.

Since the *Gr8a* mutation is not spatially restricted in *Gr8a* mutant males, it is possible that at least some of the effects of the *Gr8a* mutation on the pheromone profiles of males are indirectly mediated via its action in pheromone-sensing GRNs, instead of directly mediated via its action in oenocytes. Therefore, we next examined the effect of oenocyte-specific *Gr8a* knockdown on the production of male CHCs. We found that oenocyte-specific *Gr8a* RNAi knockdown in males leads to significant changes in their overall CHC profile relative to control males (Figure 3G). In contrast, fat-body-specific knockdown of *Gr8a* has no effect on the CHC profiles of males (Figure 3H). These data suggest that *Gr8a* is likely to play an oenocyte-specific role in the production of male CHCs. Together, our behavioral and pheromonal data indicate that *Gr8a* action contributes to mating decisions in females by co-regulating the perception of an inhibitory mating pheromone by females and males, as well as its production by males. This is consistent with a pleiotropic function for *Gr8a.*

### Gr8a-associated CHCs inhibit normal courtship behaviors

To further characterize whether any of the individual CHCs regulated by *Gr8a* actually function as inhibitory mating pheromones, we tested the effect of perfuming naïve males with individual candidate CHCs on the copulation latency of wild-type females (Ben-Shahar et al., 2010; Ben-Shahar et al., 2007; Leitner and Ben-Shahar, 2020; Lu *et al*., 2012; Lu *et al*., 2014). We found that wild-type females did not copulate with *Gr8a* mutant males that were perfumed with the alkenes 9-C_25_, 7-C_25_, and 7-C_27_ (Figure 4A). Similarly, we found that wild-type males exhibited a longer courtship latency and lower courtship index towards wild-type females perfumed with 9-C_25_ (Figure 4B-D), and exhibited longer copulation latency towards wild-type females perfumed with 7-C_25_ (Figure 4E-G). In contrast, perfuming wild-type females with 7-C_27_ had no effect on male courtship or female mating latency (Figure 4H-J). These data suggest that at least some of the CHCs regulated by *Gr8a* activity in the male oenocytes are inhibitory mating pheromones.

**Figure 4.**
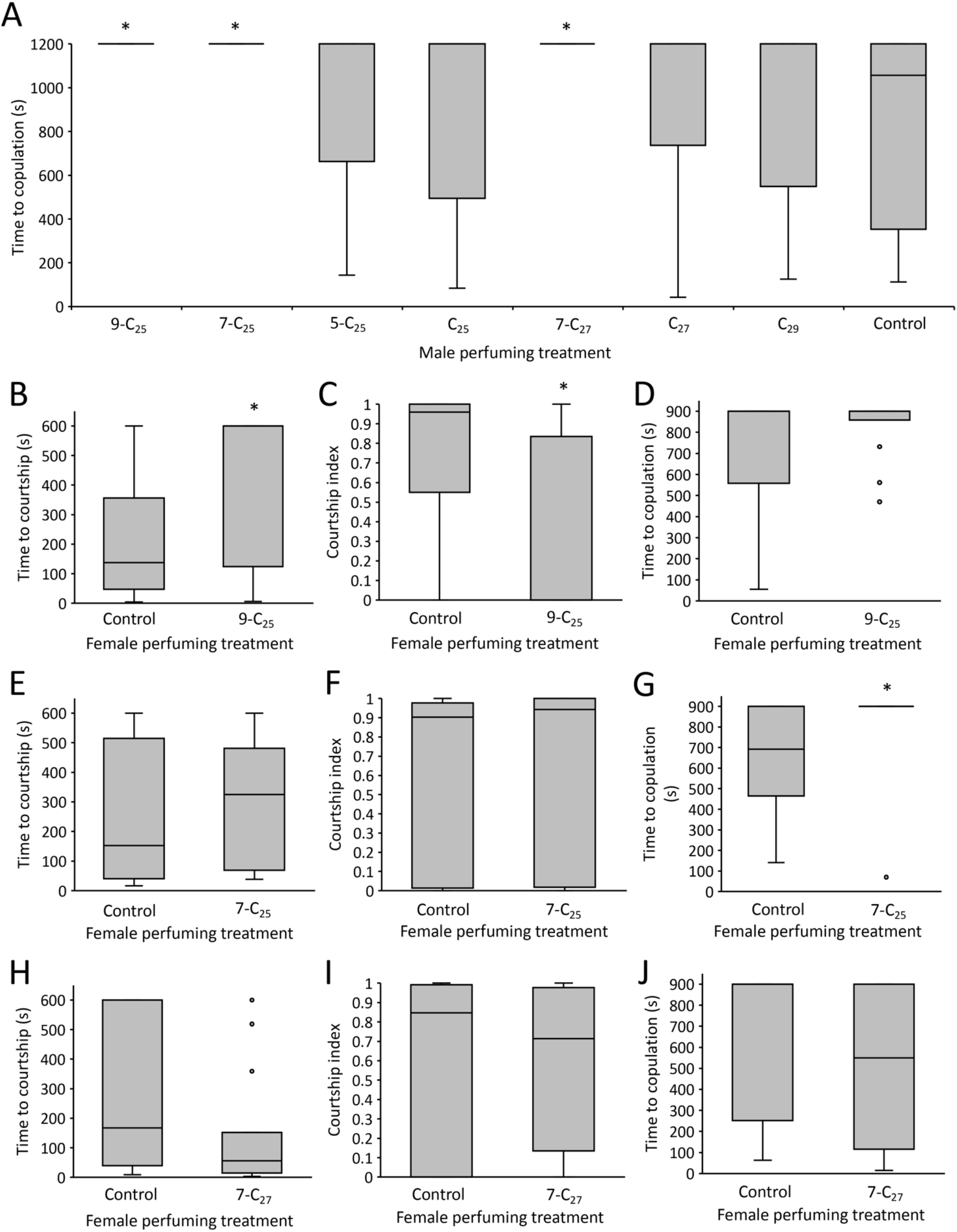
*Gr8a-*associated alkenes inhibit normal courtship behaviors. **(A)** Perfuming males with exaggerated amounts of several alkenes increases copulation latency compared to control males. **(B-D)** Perfuming females with 9-C25 increases courtship latency (B), decreases courtship index (C), but does not affect copulation latency (D) compared to control females. **(E-G)** Perfuming females with 7-C25 does not affect courtship latency (E) or index (F), but increases copulation latency (G) compared to control females. **(H-J)** Perfuming females with 7-C27 does not affect courtship latency (H), courtship index (I), or copulation latency (J) compared to control females. Asterisks above bars indicate statistically significant contrasts compared to control flies, p<0.05, Kruskal-Wallis Test followed by Dunn’s Test (A) or Mann Whitney Rank Sum Test (B-J), n=15/group.

### Variations in Gr8a contribute to species-specific male pheromonal profiles across the Drosophila genus

As populations diversify, pheromonal signals and their receptors often have to co-evolve to maintain behavioral species boundaries (Boake, 1991; Khallaf et al., 2021; Symonds and Elgar, 2008; Symonds and Wertheim, 2005). One possible mechanism for maintaining the functional coupling of coevolving signal-receptor pairs during speciation is pleiotropy (Boake, 1991; Shaw *et al.*, 2011; Singh and Shaw, 2012). Because our data suggest that *Gr8a* is a pleiotropic pheromone receptor, we tested the hypothesis that cross-species variations in the *Gr8a* coding sequence may have contributed to the rapid evolution of mating pheromones in the *Drosophila* species group (Khallaf *et al*., 2021; Shahandeh et al., 2018; Shirangi *et al*., 2009). To test this hypothesis, we first performed a phylogenetic analysis of *Gr8a* orthologs across *Drosophila* species, which indicated that *Gr8a* is a conserved, sexually dimorphic receptor across the *Drosophila* genus (Figure 5A-B). Furthermore, alignment of *Gr8a* proteins across all the major *Drosophila* clades revealed that, in spite of its high overall sequence conservation, the *Gr8a* receptor has at least one phylogenetically variable domain (magenta frame, Figure 5C), which includes the second intracellular and extracellular domains (Figure 5D).

**Figure 5.**
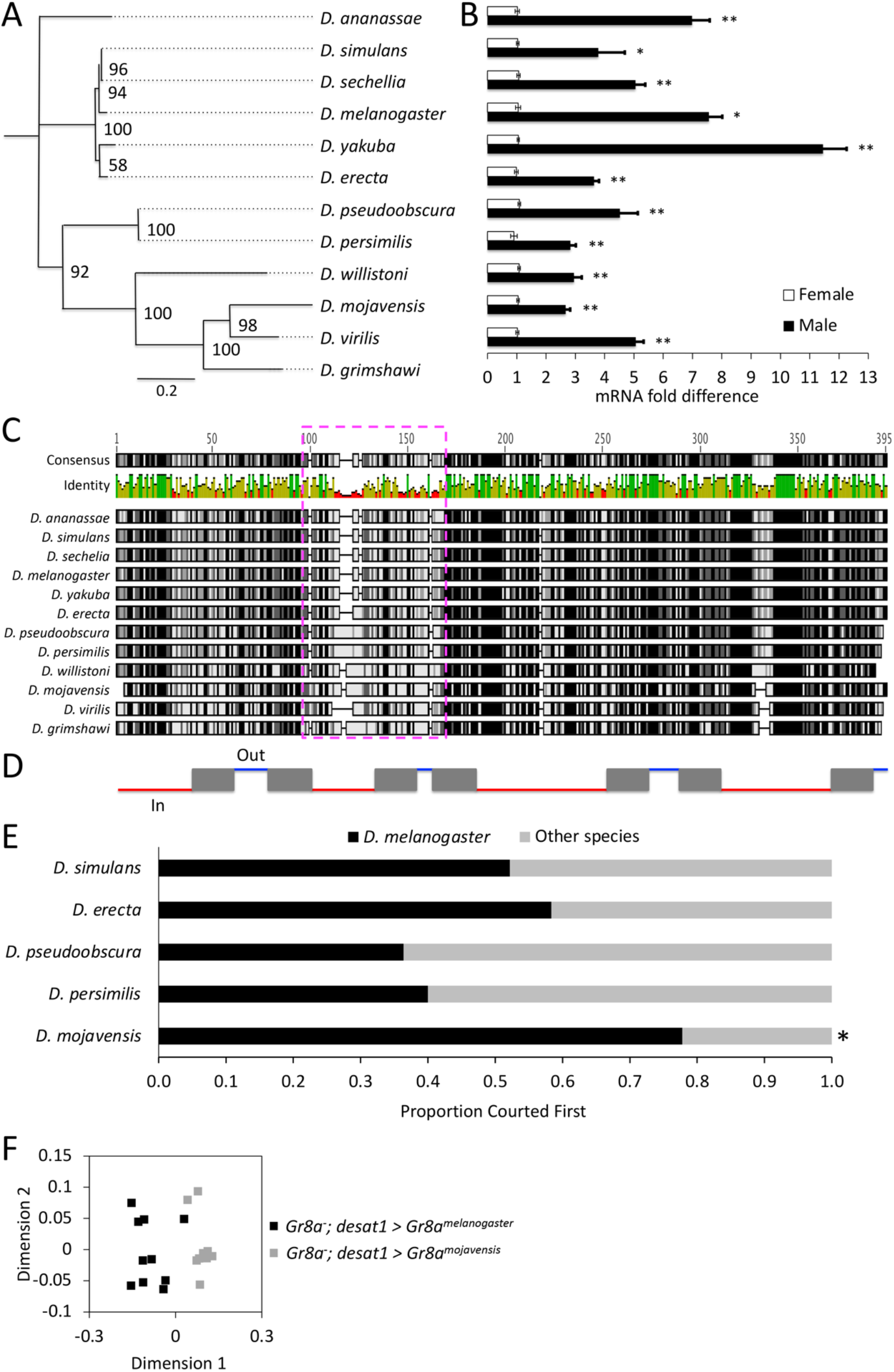
Sexually dimorphic *Gr8a* expression across the *Drosophila* genus may contribute to species-specific differences in male CHC profiles. **(A)** Phylogenetic tree of *Drosophila Gr8a* proteins. Substitution rate = 0.2. **(B)** *Gr8a* mRNA expression is enriched in males relative to females across *Drosophila*. Black, males; white, females. *, p<0.05; **,p<0.01; Mann Whitney Rank Sum Test, n=4/group. Live *D. grimshawi* was not analyzed because live specimens were not available at the *Drosophila* Species Stock Center (DSSC). **(C)** Multiple aligned amino acid sequences of *Gr8a* protein sequences from 12 species across *Drosophila.* The magenta dashed box highlights a putative hypervariable protein domain. Numbers on top of alignment indicate amino acid number. Black, 100% identical; Dark Gray, 80-100% similar; Light Gray, 60-80% similar; White, less than 60% similar (Blosum62 score matrix, threshold=1). Bars below consensus represent overall level of amino acid conservation. **(D)** *Gr8a* protein topology. Boxes, transmembrane domains; Red lines, intracellular domain; Blue lines, extracellular domains. **(E)** In female choice assays, *D. melanogaster* males court females from most other *Drosophila* species first at an equal proportion as *D. melanogaster* females, but court *D. mojavensis* females first at a lower proportion than *D. melanogaster* females. Assays performed under red light. *, p < 0.05, Pearson’s Chi-squared test. **(F)** *Gr8a* mutant *D. melanogaster* males with oenocyte-specific *D. melanogaster Gr8a* rescue differ in CHC profile from *Gr8a* mutant *D. melanogaster* males with oenocyte-specific *D. mojavensis Gr8a* rescue. Depicted as NMDS plot with Bray-Curtis dissimilarity; *Gr8a^-^; desat1 > Gr8a^melanogaster^, D. melanogaster Gr8a* oenocyte rescue; *Gr8a^-^; desat1 > Gr8a^mojavensis^. D. mojavensis Gr8a* oenocyte rescue. Bold letters in legend denote statistical significance, p < 0.05, permutation MANOVA.

Although the ligand-binding domains of the insect *Gr* gene family have not been identified yet, these data suggest that this phylogenetically variable protein domain in *Gr8a* may contribute to species-specific shifts in ligand-binding specificity and/ or sensitivity across the *Drosophila* genus. Therefore, we next tested whether the transgenic rescue of the *Gr8a* null allele via ectopic expression of *Gr8a* cDNAs from different *Drosophila* species is sufficient to drive changes in the CHC profile of *D. melanogaster* males. By using a cross-species male mate-choice assay, we found that while *D. melanogaster* males are generally promiscuous, they do court *D. mojavensis* females at a significantly lower proportion than conspecific females. Because these assays are performed under red light, which eliminates visual mating cues, these data suggested that the lower sex drive towards *D. mojavensis* females is pheromone-dependent (Figure 5E). Subsequently, we generated transgenic *D. melanogaster* lines which express either the *D. mojavensis* or the *D. melanogaster Gr8a* cDNAs driven by an oenocyte-specific GAL4 in the background of the *Gr8a* null allele. Comparison of male CHC profiles across the two genotypes revealed that rescuing the *Gr8a* mutation by *Gr8a* cDNAs from these two distantly related species resulted in significantly different male CHC profiles (Figure 5F). These data indicate that species-specific *Gr8a* coding variations are sufficient to drive differential CHC production by the male oenocytes, and suggest that pleiotropic pheromone receptors may have played a role in driving the rapidly evolving behavioral mating boundaries in *Drosophila.*

## DISCUSSION

The data presented here demonstrate that *Gr8a* is a pleiotropic chemoreceptor that coregulates the perception and production of an inhibitory pheromonal signal that plays an important role in mating behaviors of both *D. melanogaster* sexes. How *Gr8a,* a member of a canonical chemoreceptor family, might also contribute to the production of pheromonal signals is not obvious. In some better understood secretory cell types, autoreceptors are essential for the regulation of synthesis and secretion rates. For example, dopaminergic and serotonergic cells regulate rates of synthesis and release of their respective neuromodulators by the action of autoreceptors, which act via signaling feedback in response to changes in the extracellular concentrations of the secreted molecule (Ford, 2014; Stagkourakis et al., 2016). Therefore, one possible explanation for how *Gr8a* might regulate the synthesis and/or secretion of specific CHCs is by acting as an oenocyte-intrinsic autoreceptor, which regulates the synthesis of specific CHCs by providing feedback information about their levels in internal stores and/ or extracellularly (Figure 6).

**Figure 6.**
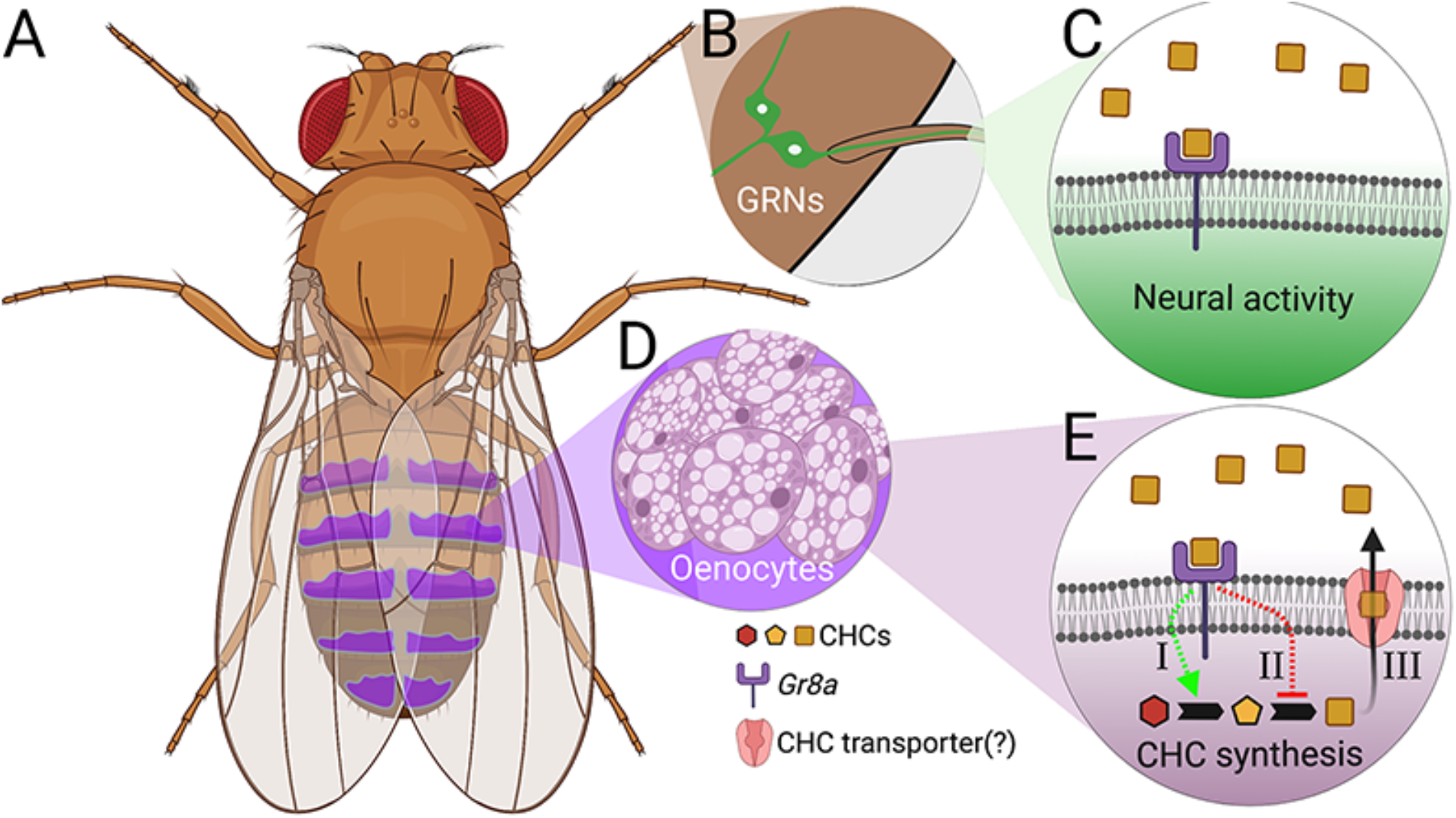
Model for the pleiotropic action of *Gr8a* in the perception and production of pheromones. **(A)** *Drosophila* male. The location of CHC-producing oenocytes is shown in magenta. **(B)** *Gr8a*-expressing GRNs are located at the last tarsal segment of the prothoracic legs. **(C)** *Gr8a* functions as an inhibitory pheromone receptor in a specific subset of leg GRNs. **(D)** Oenocytes are the primary CHC-producing cells in the male abdomen. **(E)** *Gr8a* functions as an autoreceptor in oenocytes, which regulates CHC synthesis [I-II] and/or CHC secretion [III] via signaling feedback loops.

Recent studies have indicated that *Drosophila* bitter receptor neurons typically express multiple *Gr* genes, and that bitter receptor ligand specificity is determined via combinatorial heteromeric receptor complexes (Dweck and Carlson, 2020; Shim *et al*., 2015; Sung et al., 2017). *Gr8a* is specifically required for the sensory perception of the feeding deterrent L-canavanine (Lee *et al.*, 2012; Shim *et al.*, 2015), but not for the detection of other bitter feeding deterrents such as caffeine, strychnine, and umbelliferone (Lee et al., 2009; Poudel et al., 2015). Our data indicate that similar to other *Drosophila* “bitter” taste receptors (Lacaille *et al*., 2007; Moon *et al*., 2009), *Gr8a* contributes to inhibitory sensory inputs in the contexts of both feeding and mating decisions. In the context of feeding, *Gr8a*-dependent perception of L-canavanine is mediated via its heterotrimeric interaction with *Gr66a* and *Gr98b* in bitter sensing neurons in the proboscis (Shim *et al*., 2015). However, although both *Gr66a* and *Gr98b* were also identified in our initial screen for receptors enriched in the adult abdomen, we found that *Gr66a* is expressed in both sexes and *Gr98b* is specifically enriched in females (Table 1). Therefore, we conclude that *Gr8a*-dependent contributions to sensory functions associated with mating decisions are independently driven via its heteromerization with different *Gr* genes than those that drive feeding-specific decisions.

Although we do not yet know the specific chemical identity of the ligand of *Gr8a,* previous studies indicated that at least two inhibitory mating pheromones, 11-cis-vaccenyl acetate (cVA) and CH503, are transferred from males to females during copulation. While our data suggest that the *Gr8a* mutation affects the level of cVA expressed by males, it is unlikely that either cVA or CH503 are the putative *Gr8a* ligands because the volatile cVA acts primarily via the olfactory receptor *Or67d* (Benton *et al*., 2007; Datta *et al*., 2008; Kurtovic *et al*., 2007), and CH503 has been reported to signal via *Gr68a*-expressing neurons, which are anatomically distinct from the *Gr8a* GRNs we describe here (Figure 1A-B) (Shankar et al., 2015; Yew et al., 2009). Instead, our analyses of the effect of the *Gr8a* mutation on the CHC profile (Figure 3), and our results of the perfuming behavioral studies (Figure 4), suggest that the alkenes 5-C_25_, 7C_25_, and 7-C_27_, which seem to act as inhibitory mating signals as well, are potentially the ligands of *Gr8a.*

Overall, our studies indicate that pleiotropic receptors, such as *Gr8a,* contribute to the physiological coupling between the production and perception of some mating pheromones by acting as both a sensory receptor in pheromone-sensing neurons, and possibly as an autorecepor for the same chemical in the pheromone-producing oenocytes. Our finding that *Gr8a* is also a sexually dimorphic receptor that is conserved across the *Drosophila* genus, with at least one phylogenetically variable domain (Figure 5A-C), suggests that it might also drive the divergence of mating signaling systems in association with rapid speciation. This is supported by our finding that rescuing the *Gr8a* mutation in *D. melanogaster* with a *Gr8a* cDNA from a distant species, *D. mojavensis,* leads to the development of a male CHC profile that is different from the profile of mutant males rescued with the *D. melanogaster Gr8a* cDNA (Figure 5F).

Studies in other animal species suggest that receptor pleiotropy likely plays a role in mating signaling via other sensory modalities including auditory communication in crickets (Heinen-Kay et al., 2020; Hoy *et al*., 1977; Wiley *et al*., 2012) and visual communication in fish (Fukamachi et al., 2009). While the specific genes and signaling pathways that mediate the coupling of the mating signals and their receptors in these mating systems remain mostly unknown, these data suggest that genetic linkage in signal-receptor pairs important for mating communication is likely to be more common than previously thought. Therefore, the genetic tractability of *D. melanogaster,* in combination with the diversity of mating communication systems in this species-rich phylogenetic group, provide a unique opportunity for understanding the evolution and mechanisms that drive and maintain the robustness of mating systems at the genetic, molecular, and cellular levels.

## ACKNOWLEDGMENTS

We thank members of the Ben-Shahar lab for comments on earlier versions of the manuscript. We thank Joshua Krupp (University of Toronto) for assistance with perfuming studies, Nabeel Chowdhury and Deanna Simon for assistance with qRT-PCR analysis, and Paula Kiefel for technical help with generating transgenic flies. This work was supported by NSF grants 1322783, 1754264, and 1707221, and NIH grant NS089834 awarded to Y. B-S. Stocks obtained from the Bloomington Drosophila Stock Center (NIH P40OD018537) were used in this study. Wild-type *Drosophila* species were obtained from the National *Drosophila* Species Stock Center at Cornell University.

## AUTHOR CONTRIBUTIONS

K.M.Z., C.V., J.G.M. and Y.B-S designed experiments. K.M.Z., C.V., N.L., X.L., S.H., J.G.M. and Y.B-S collected and analyzed data. K.M.Z., C.V., N.L. and Y.B-S wrote the manuscript.

## METHODS

### Animals

Flies were maintained on a standard cornmeal medium under a 12:12 light-dark cycle at 25 Celsius. Unless specifically stated, the *D. melanogaster Canton-S (CS)* strain served as wild-type control animals. UAS-TNT-E, UAS-TNT-IMP-V1-A, UAS-mCD8::GFP, UAS-myr::GFP, UAS-Red Stinger, Df(1)BSC663, Df(1)BSC754, *Gr8a-GAL4, Gr8a^1^, desat1* -Gal4, r4-Gal4 and *fruP1-LexA* fly lines were from the Bloomington Stock center. Originally in the *w^1118^* background, the *Gr8a^1^ null* allele was outcrossed for six generations into the *CS* wild-type background, which was used as a control. Likewise, the *desat1-Gal4* allele was outcrossed for six generations into this *Gr8a null* background. *PromE(800)-GAL4* and *PromE(800)>Luciferase* were from Joel Levine (The University of Toronto, Canada). The following *Drosophila* species were obtained from the San Diego Stock Center: *D. simulans* 14011-0251.192, *D. sechellia* 14021-0248.03, *D. yakuba* 14021-0261.01, *D. erecta* 14021-0224.00, *D. ananassae* 14024-0371.16, *D. pseudoobscura* 14011-0121.104, D. *persimilis* 14011-0111.50, *D. willistoni* 14030-0811.35, *D. mojavensis* 15081-1352.23, and *D. virilis* 15010-1051.118. The UAS-*Gr8a* transgenic lines were generated by cloning the *D. melanogaster* and *D. mojavensis Gr8a* cDNAs into pUAST-attB vector by using 5’ EcoRI and 3’ NotI restriction sites, followed by *ΦC31* integrase-dependent transgenesis at a Chromosome 2 *attP* landing site (2L:1476459), as previously described (Zheng et al., 2014). Subsequently, both *UAS-Gr8a^CDNA^* lines were transgressed into the *Gr8a^1^* background, resulting in complete substitution of the endogenous *Gr8a* with expression of a *Gr8a* ortholog. The *ppk23*-LexA line was generated by integrating our previously described *ppk23* promotor DNA fragment (Lu *et al*., 2012) into the pBPnlsLexA::p65Uw plasmid (Pfeiffer et al., 2010), followed by *ΦC31* integrase-dependent transgenesis as above.

The GFP-tagged allele of *Gr8a* was generated via CRISPR/*Cas9*-dependent editing using a modified “scarless” strategy by using the sgRNA CGAGCAAGGCGGGAACGATT and a 3XP3>dsRed in the donor plasmid as a reporter for edited animals as previously described (Hill et al., 2017; Hill et al., 2019). Control lines with matching genetic backgrounds were established by selecting DsRed-negative injected animals. The final tagged *Gr8a* allele was generated by removing the DsRed cassette via the introduction of the *piggyBac* transposase (Hill *et al*., 2019).

### Immunohistochemistry

To visualize the expression pattern of *Gr8a* in males and females, *Gr8a-GAL4* flies (Lee et al., 2012) were crossed to UAS-CD8::EGFP and live-imaged at 5 days old using a Nikon-A1 confocal microscope. To demonstrate *Gr8a* expression in oenocytes, abdomens from *Gr8a-GAL4/UAS-myr::GFP; PromE(800)>Luciferase* flies were dissected and immunostained as previously described (Lu et al., 2012; Zheng et al., 2014) by using a Rabbit anti-GFP (1:1000; A-11122, Thermo Fisher Scientific) and a mouse anti-luciferase (1:100; 35-6700, Thermo Fisher Scientific) antibodies followed by AlexaFluor 488 anti-rabbit and AlexaFluor 568 anti-mouse secondary antibodies (Both at 1:1000; Thermo Fisher Scientific). To visualize the GR8A protein, abdomens of control flies and flies with CRISPR/Cas9 generated GFP-tagged GR8A were dissected and immunostained as previously described (Lu et al., 2012; Zheng et al., 2014) using a Rabbit anti-GFP antibody (1:1000; A-11122, Thermo Fisher Scientific) followed by AlexaFluor 488 anti-rabbit secondary antibody (1:1000; Thermo Fisher Scientific).

### mRNA expression

Newly eclosed flies were separated by sex under CO2 and aged for 5 days on standard cornmeal medium. On day 6, flies were placed in a −80°C freezer until RNA extraction. To separate body parts, frozen flies were placed in 1.5ml microcentrifuge tubes, dipped in liquid nitrogen, and then vortexed repeatedly until heads, appendages, and bodies were clearly separated. Total RNA was extracted using the Trizol Reagent (Thermo Fisher Scientific) separately from heads, bodies, and appendages for *Gr8a* expression and from bodies for desaturase enzyme genes. cDNAs were synthesized using SuperScript II reverse transcriptase (Thermo Fisher Scientific) with 500 ng total RNA in a 20 uL reaction. Real-time quantitative RT-PCR was carried out as previously described with *Rp49* as the loading control gene (Hill *et al*., 2017; Hill *et al*., 2019; Lu *et al*., 2012; Lu *et al*., 2014; Zheng *et al*., 2014). Primer sequences are described in Supplemental Tables 1–3.

### Courtship Behavior Assays

Single-pair assays were performed as we have previously published (Lu et al., 2012, 2014). In short, newly eclosed males were kept individually on standard fly food in plastic vials (12 x 75mm). Newly eclosed virgin females were kept in groups of 10 flies. All behaviors were done with 4-7 day-old animals, which were housed under constant conditions of 25° C and a 12h:12h light-dark cycle. Courtship was video recorded for 10 min for male courtship and 15 min for female mating receptivity. Male courtship latency and index were measured as previously described (Lu *et al*., 2012; Lu *et al*., 2014). Female receptivity index was defined as the time from the initiation of male courtship until copulation was observed. Unless otherwise indicated, assays were performed under normal light conditions.

Male mate-choice assays were performed in round courtship arenas. Briefly, one *D. melanogaster* virgin female and one interspecific virgin female was decapitated under CO2 and placed in the arena. One virgin male *D. melanogaster* was then aspirated into the arena and behavior was video recorded for 10 minutes. The first female courted (by male wing extension) was noted. Male mate-choice assays were performed under red light conditions.

### Perfuming studies

Synthetic compounds were synthesized by J.G.M. Perfuming studies were performed using a modified protocol from (Billeter et al., 2009). In short, 3 mg of each compound was dissolved in 6 mL hexane (Sigma-Aldrich #139386-500ML) and 0.5 mL was pipetted into individual 2 mL glass vials fitted with 9mm PTFE lined caps (Agilent Crosslab, Santa Clara, CA, USA). The hexane was evaporated under a nitrogen gas flow, such that a residue of the compound was left around the bottom one-third of the vial. Control vials were prepared using hexane without a spiked compound. Vials were kept at −20°C until use. Flies used in these trials were collected as described above, kept in single sex groups and aged for 4 days on standard cornmeal medium at 25°C. 24 hours before perfuming, 20 flies of one or the other sex were placed in glass vials containing standard cornmeal medium (12 x 75mm). To perfume the flies, these groups of 20 flies were dumped without anesthesia into each 2 mL vial containing the compound of interest, and were vortexed at medium-low speed for 3 pulses of 20 seconds punctuated by 20 second rest periods. Flies were transferred to new food vials and were allowed to recover for one hour. Perfumed flies were then used in courtship behavior assays as described above and the remaining flies were used in pheromone analyses to verify compound transfer. The genotype of flies that were perfumed differed based upon the genotype with the lower amount of each compound as determined in Figure 3 (B, C, F). In all cases, compound transfer was verified by CHC extraction and GC/MS (Supplemental Table 4).

### Phylogenetic analysis

Protein sequences of GR8A orthologs from the 12 sequenced *Drosophila* reference genomes were aligned by using the ClustalW algorithm in the Omega package (Sievers et al., 2011), followed by ProtTest (v2.4) to determine the best model of protein evolution (Abascal et al., 2005). Subsequently, Akaike and Bayesian information criterion scores were used to select the appropriate substitution matrix. We then used a maximum likelihood approach and rapid bootstrapping within RAxML v 7.2.8 Black Box on the Cipres web portal to make a phylogenetic tree (Miller et al., 2010). Visualizations of the bipartition files were made using FigTree v1.3.1 (http://tree.bio.ed.ac.uk/software/figtree/).

### Pheromone Analysis

Virgin flies were collected upon eclosion under a light CO^2^ anesthesia and kept in single-sex vials in groups of 10 with 6 biological replications for each genotype and sex. Virgin flies were aged for 5 days on standard cornmeal medium at 25°C. To collect mated flies, both females and males were aged for 3 days before single mating pairs were placed in a standard fly vial with standard cornmeal food for 24 hours. The pair was then separated for 24 hours before collection. Copulation was confirmed by the presence of larvae in the vials of mated females several days later. On the morning of day 5, flies were anesthetized under light CO^2^ and groups of five flies were placed in individual scintillation vials (VWR 74504-20). To extract CHCs, each group of flies was covered by 100 uL hexane (Sigma-Aldrich #139386-500ML) containing 50μg/mL hexacosane (Sigma-Aldrich #241687-5G) and was washed for ten minutes. Subsequently, hexane washes were transferred into a new 2 ml glass vial containing a 350 uL insert (Thermo Scientific C4000-LV-1W) and were stored at −20°C until shipment to the Millar laboratory.

Analyses of CHC profiles were done by gas chromatography and mass spectroscopy (GC-MS) in the Millar laboratory at UC Riverside as previously described (Chung et al., 2014). Peak areas were measured, and data was normalized to known quantity of internal standard hexacosane (Sigma-Aldrich #241687-5G). The relative proportion of each compound in each sample was calculated and used in further statistical analysis.

### Statistical Analysis

All statistical analyses were performed in R (v 3.6.2). The following functions were used in the base statistics package: t.test() (t-test), wilcox.test() (Mann-Whitney Rank Sum Test), aov() (ANOVA), TukeyHSD() (Tukey’s HSD post hoc test), Kruskal.test() (Kruskal-Wallis test), chisq.test() (Pearson’s Chi-squared test). Kruskal-Wallis post hoc was performed using the dunn.test.control function in the PMCMR package (Pohlert, 2014). Qualitative CHC data were analyzed through a permutation MANOVA using the adonis function in the vegan package of R with Bray-Curtis dissimilarity measures (Oksanen, 2011). CHC profile data were visualized using non-metric multidimensional scaling (metaMDS) function in the vegan package of R (Oksanen, 2015) using Bray-Curtis dissimilarity, and either 2 or 3 dimensions in order to minimize stress to <0.1.

## SUPPLEMENTAL INFORMATION

**Supplemental Figure 1.**
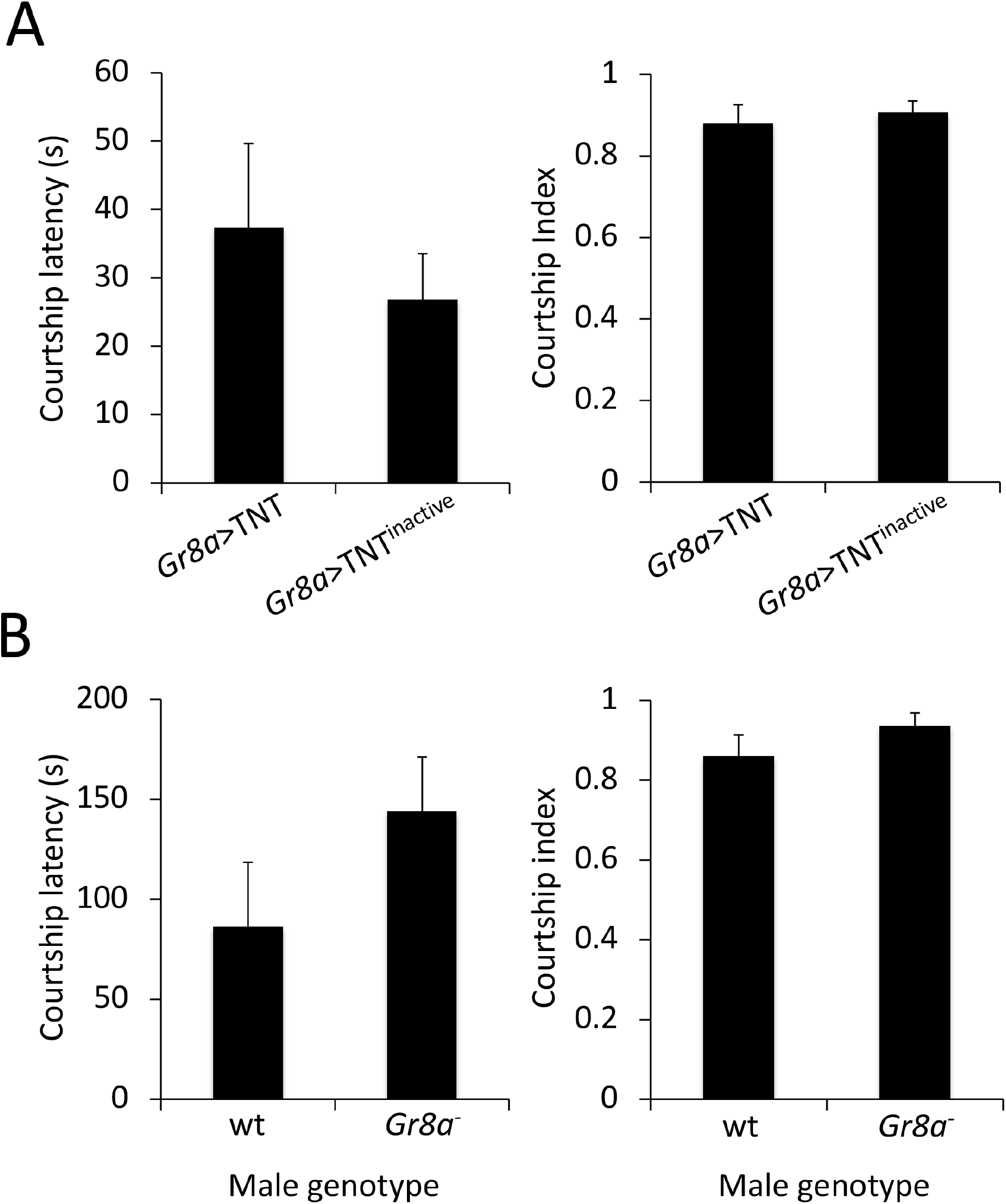
*Gr8a* has no effect on male courtship latency or index toward wild-type females. (A) Courtship latency (s) and index of control *Gr8a-gal4/UAS-IMP-TNT-V1A* (*Gr8a*>TNT^inactive^) and *Gr8a-gal4/UAS-TNT-E (Gr8a>TNT)* mutant males towards wild-type females. (B) Courtship latency (s) and index of wild-type (CS) and *Gr8a null* (Gr8a^-^) males toward wild-type decapitated females. Mann Whitney Rank Sum Test, not significant (p>0.05), n=15/group.

**Supplemental Table 1.**
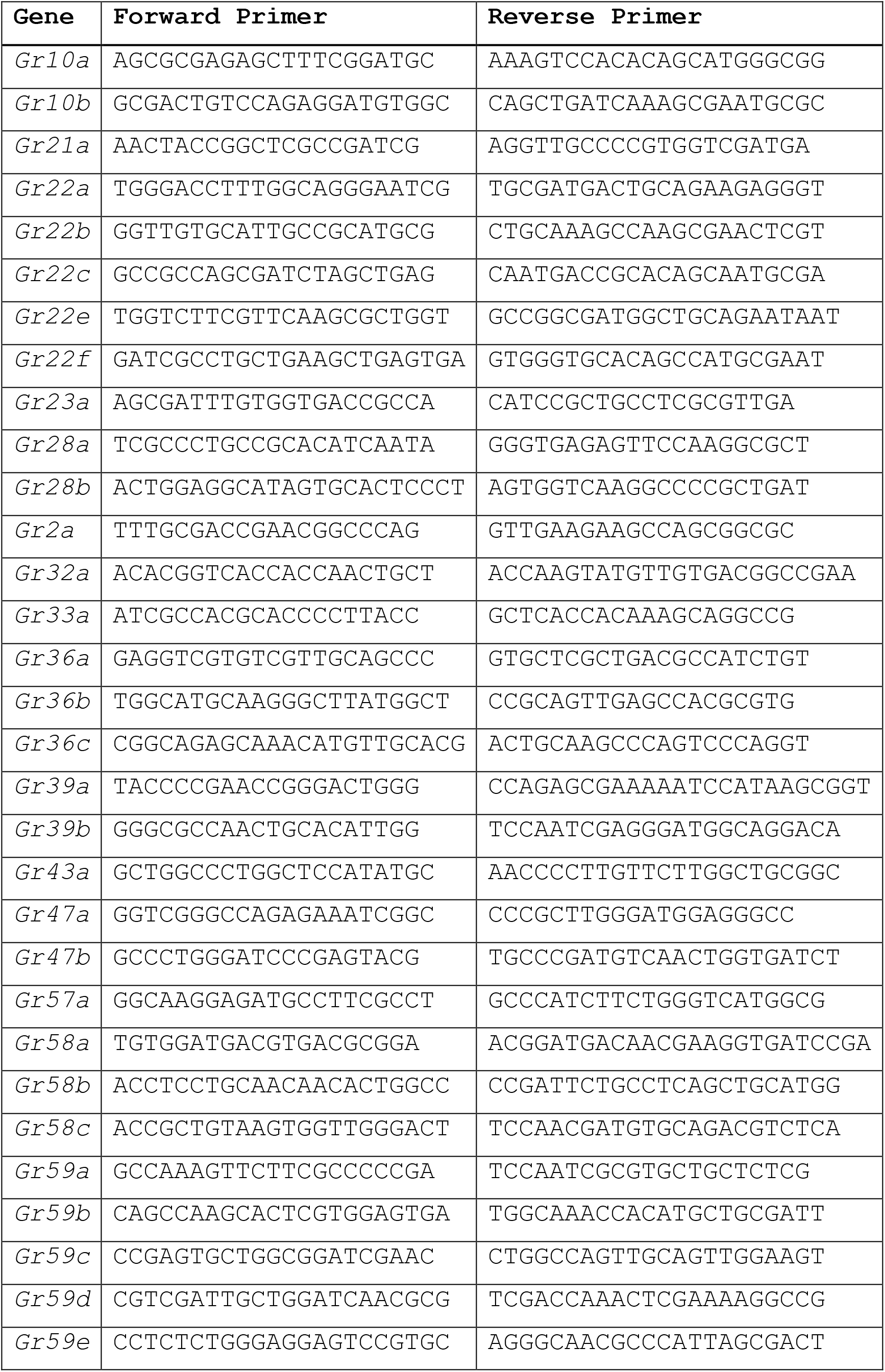
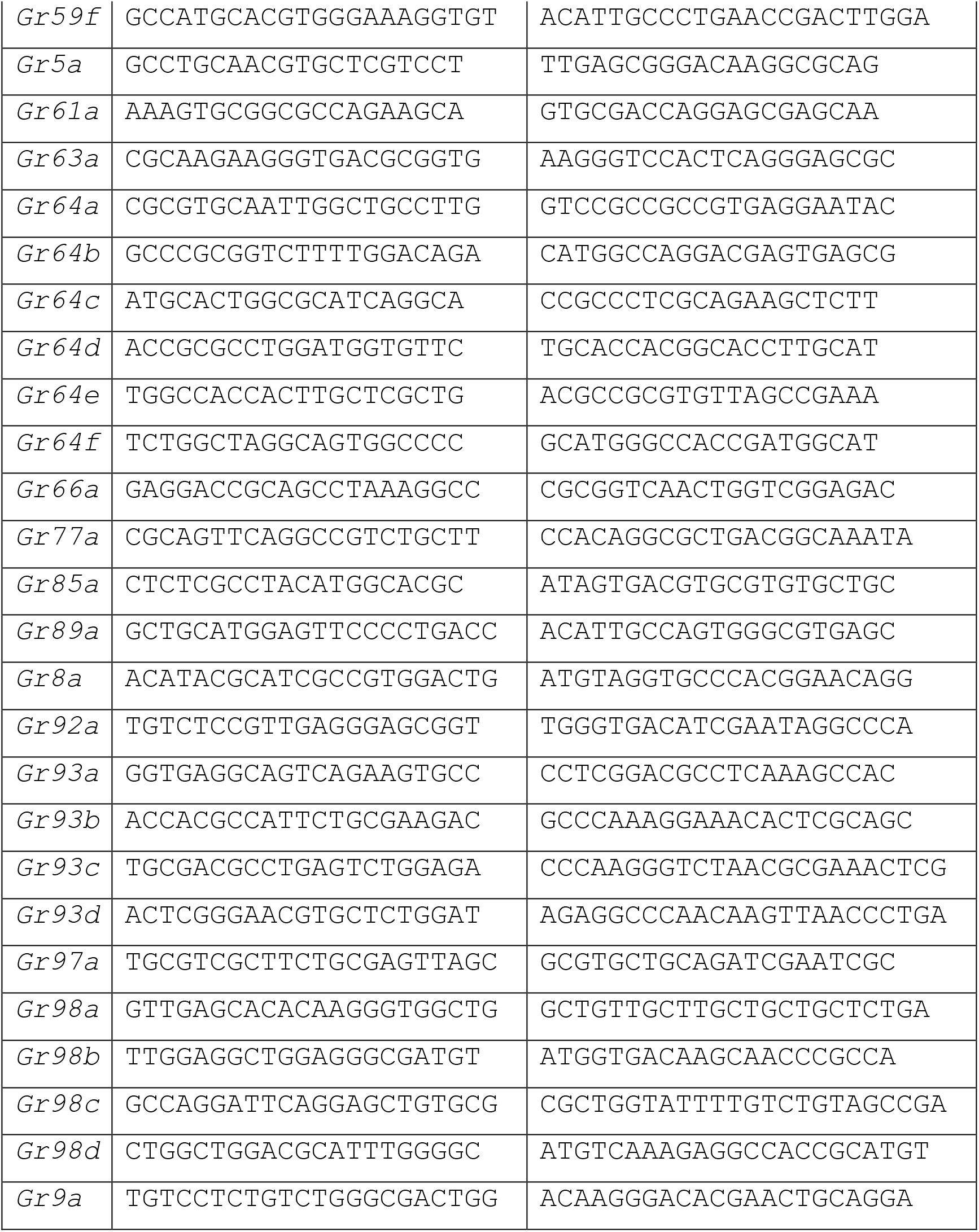
Nucleotide sequences for qRT-PCR primers for *D. melanogaster Gr* genes.

**Supplemental Table 2.**
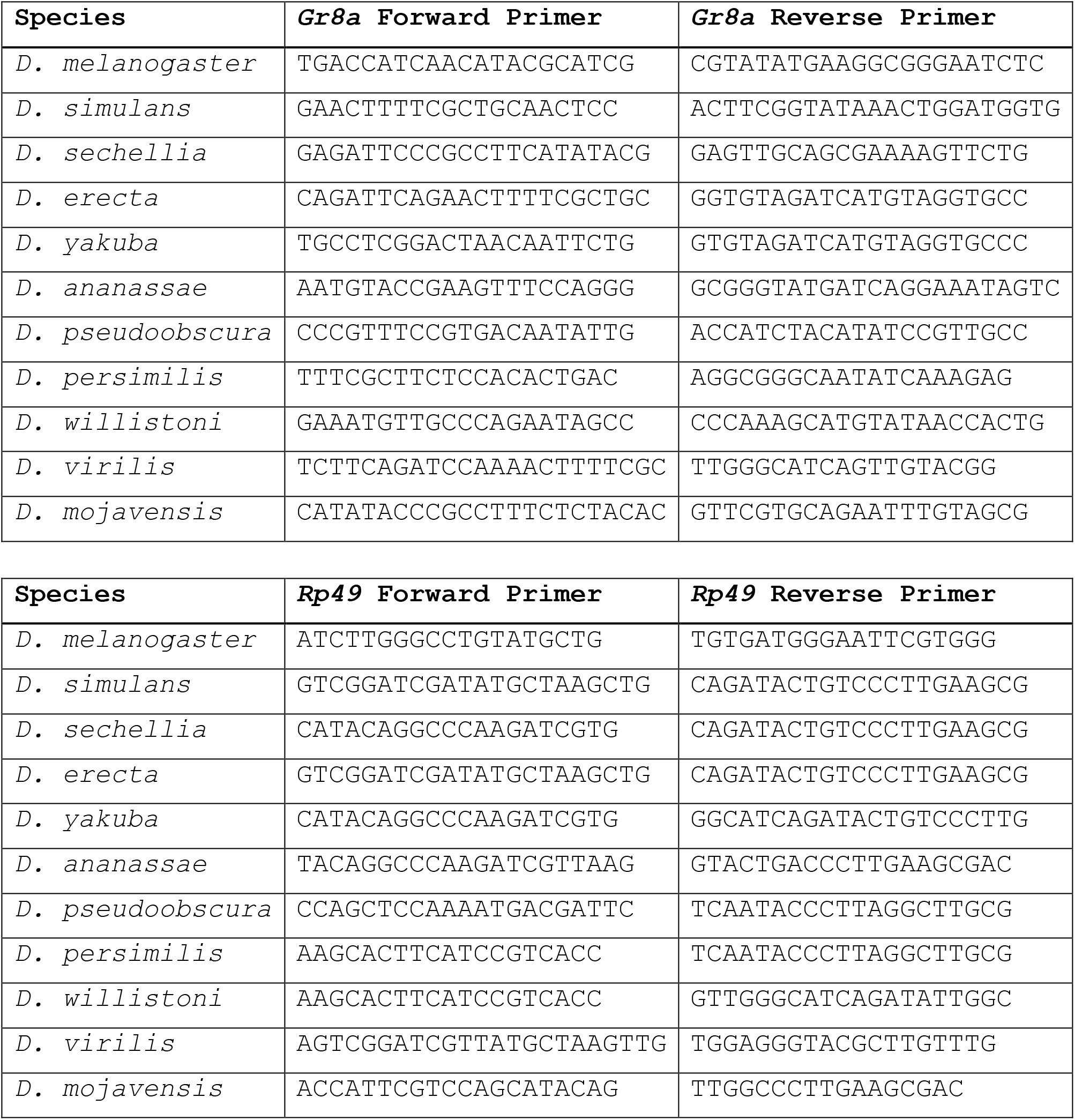
Nucleotide sequences for qRT-PCR primers for *D. melanogaster Gr8a, Rp49* and orthologs.

**Supplemental Table 3.**
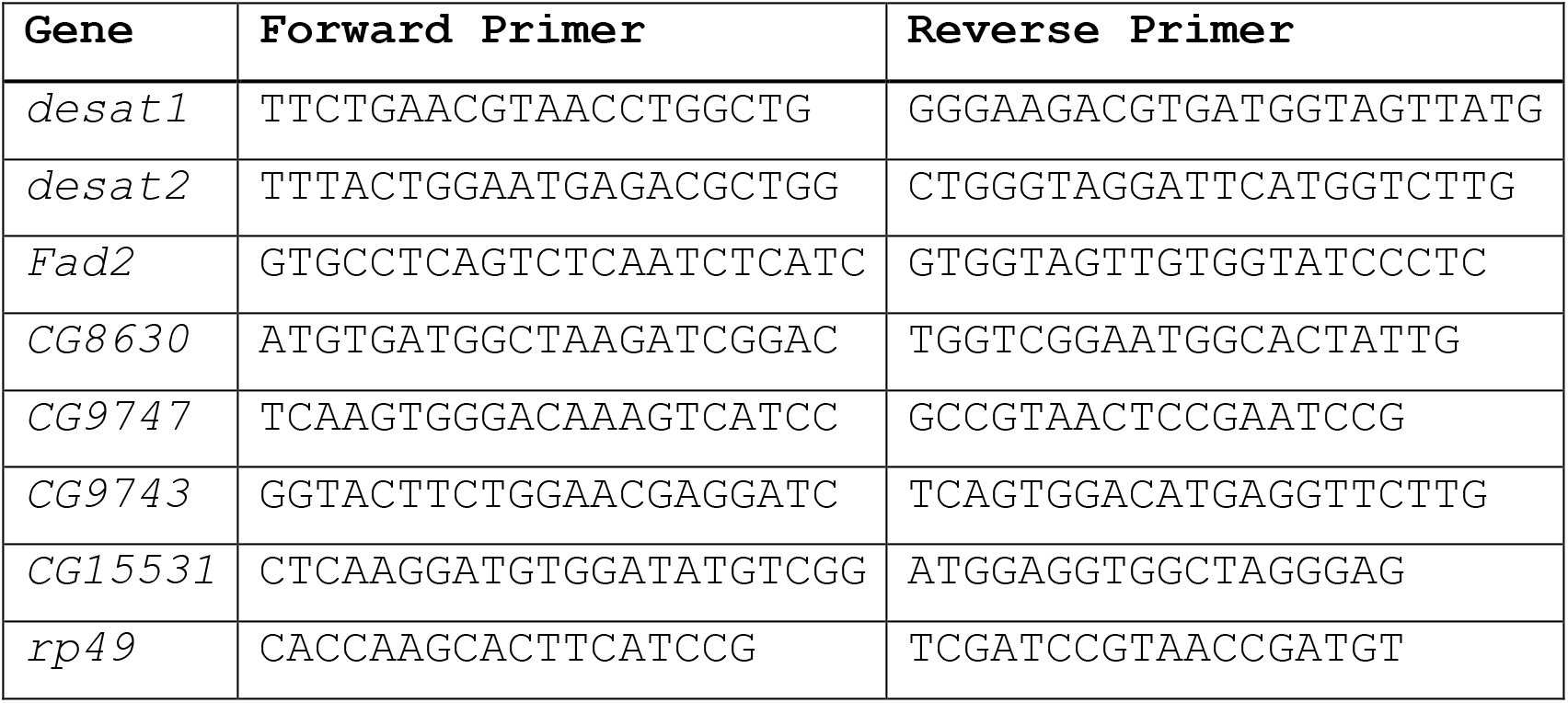
Nucleotide sequences for qRT-PCR primers for *D. melanogaster* desaturase enzyme genes.

**Supplemental Table 4.** Amount (ng) of each perfumed compound measured in each sample of perfumed flies (5 flies per sample).

| Compound | Genotype | Sex | ng/sample | ng/fly |
| --- | --- | --- | --- | --- |
| C<sub>25</sub> | wt | male | 1350 | 270 |
| C<sub>25</sub> | wt | male | 1170 | 234 |
| C<sub>25</sub> | wt | male | 27730 | 5546 |
| 9-C<sub>25</sub> | Gr8a- | male | 67390 | 13478 |
| 9-C<sub>25</sub> | Gr8a- | male | 59380 | 11876 |
| 9-C<sub>25</sub> | Gr8a- | male | 1000 | 200 |
| 7-C<sub>25</sub> | Gr8a- | male | 43790 | 8758 |
| 7-C<sub>25</sub> | Gr8a- | male | 31330 | 6266 |
| 7-C<sub>25</sub> | Gr8a- | male | 24130 | 4826 |
| 5-C<sub>25</sub> | Gr8a- | male | 12050 | 2410 |
| 5-C<sub>25</sub> | Gr8a- | male | 14160 | 2832 |
| 5-C<sub>25</sub> | Gr8a- | male | 1870 | 374 |
| C<sub>27</sub> | Gr8a- | male | 890 | 178 |
| C<sub>27</sub> | Gr8a- | male | 720 | 144 |
| C<sub>27</sub> | Gr8a- | male | 263 | 52.6 |
| 7-C<sub>27</sub> | Gr8a- | male | 36710 | 7342 |
| 7-C<sub>27</sub> | Gr8a- | male | 21250 | 4250 |
| 7-C<sub>27</sub> | Gr8a- | male | 15910 | 3182 |
| C<sub>29</sub> | wt | male | 260 | 52 |
| C<sub>29</sub> | wt | male | 340 | 68 |
| C<sub>29</sub> | wt | male | 890 | 178 |
| 9-C<sub>25</sub> | wt | female | 14060 | 2812 |
| 9-C<sub>25</sub> | wt | female | 13830 | 2766 |
| 7-C<sub>25</sub> | wt | female | 23200 | 4640 |
| 7-C<sub>25</sub> | wt | female | 12780 | 2556 |
| 7-C<sub>27</sub> | wt | female | 1010 | 202 |
| 7-C<sub>27</sub> | wt | female | 1080 | 216 |

## Supplemental Data Legends

**Figure 1 Data:** Average qRT-PCR Ct scores across 3 technical replicates for each fly sample for *Gr8a* and *rp49* (control).

**Figure 2 Data:** Copulation latency (s) or courtship index of single-pair courtship trials corresponding to Figure 2.

**Figure 3 Data:** Amount (ng) of each compound extracted from each sample (5 flies/sample) in Figure 3. Average qRT-PCR Ct scores across 3 technical replicates for each fly sample for every desaturase gene measured and *rp49* (control). Genes were run on separate qPCR plates, indicated by plate number in parentheses.

**Figure 4 Data:** Copulation latency (s) of single-pair courtship trials with perfumed males. Courtship latency (s), courtship index, and copulation latency (s) of single-pair courtship trials with perfumed females.

**Figure 5 Data:** Average qRT-PCR Ct scores across 3 technical replicates for each fly sample for *Gr8a* and *rp49* (control) across *Drosophila* species. Number of flies courted first by D. melanogaster males in choice assays. Amount (ng) of each compound extracted from each sample (5 flies/sample) in Figure 5.

**Figure S1 Data:** Courtship latency (s) and index of single-pair courtship trials corresponding to Supplemental Figure 1.

